# Expression and protein sequence analyses of zebrafish *impg2a* and *impg2b*, two proteoglycans of the interphotoreceptor matrix

**DOI:** 10.1101/2021.03.09.434550

**Authors:** M.E. Castellini, G. Spagnolli, E. Biasini, S. Casarosa, A. Messina

## Abstract

Photoreceptor outer segments projecting from the surface of the neural retina toward the retinal pigment epithelium (RPE) are surrounded by a carbohydrate-rich matrix, the interphotoreceptor matrix (IPM) [1,2]. This extracellular compartment is necessary for physiological retinal function. However, specific roles for molecules characterizing the IPM have not been clearly defined [3]. Recent studies have found the presence of nonsense mutations in the interphotoreceptor matrix proteoglycan 2 (*IMPG2)* gene in patients affected by autosomal recessive Retinitis Pigmentosa (arRP) [4,5] and autosomal dominant and recessive vitelliform macular dystrophy (VMD) [6,7]. The gene encodes for a proteoglycan synthesized by photoreceptors and secreted in the IPM. However, little is known about the function and structure of this protein. We used the teleost zebrafish (*D*.*rerio*) as a model to study *IMPG2* expression both during development and in adulthood, as its retina is very similar in humans [8]. In zebrafish, there are two IMPG2 proteins, IMPG2a and IMPG2b. We generated a phylogenetic tree based on IMPG2 protein sequence similarity among different vertebrate species, showing a significant similarity despite the evolutionary distance between humans and teleosts. In fact, human IMPG2 and *D*.*rerio* IMPG2a and IMPG2b share conserved SEA and EGF-like domains. Homology models of these domains were obtained by using the iTasser server. Finally, expression analyses of *impg2a* and *impg2b* during development and in the adult fish showed expression of both mRNAs starting from 3 days post fertilization (dpf) in the outer nuclear layer of zebrafish retina that continues throughout adulthood. This data lays the groundwork for the generation of novel and most needed animal models for the study of IMPG2-related inherited retinal dystrophies.

## Introduction

The IPM is the extracellular matrix, mainly composed of proteoglycans and glycosaminoglycans, surrounding retinal photoreceptor outer segments and ellipsoids [9]. The function of the IPM in retinal function has started to be investigated only recently, as is its involvement in retinal disorders [10]. In the last years, new roles for the IPM were identified, which include intercellular communication, membrane and matrix turnover, regulation of neovascularization, cell survival, photoreceptor differentiation and maintenance, retinoid transport [3,5,11–13]. Moreover, mutations in proteins localized to the IPM such as IRBP have been shown to be involved in inherited retinal dystrophies (IRDs) [14–17]. Recent studies have reported that mutations in the *IMPG2* gene are associated with arRP [4,5] and autosomal dominant and recessive VMD [6,7] in humans. Retinitis pigmentosa (RP [MIM 268000]) is the most common IRD [18–20] involving progressive degeneration of photoreceptor cells and RPE [21,22]. Vitelliform macular dystrophy (VMD [MIM 153700]), also called Best disease, is an early-onset disorder characterized by accumulation of lipofuscin-like material within and beneath the retinal pigment epithelium together with a progressive loss of central vision [23,24]. The *IMPG2* gene encodes for the proteoglycan IMPG2, synthesized by both rods and cones and secreted in the IPM [1,2,25, 26]. Recent studies have shown progressive cone cell degeneration, increased levels of endoplasmic reticulum (ER) stress-related proteins and abnormal accumulation of the interphotoreceptor proteoglycan 1 (IMPG1) at the subretinal space, leading to reduced visual function, in *IMPG2* knockout (KO) mouse models [27,28]. The function of IMPG2 in retinal development and function, however, has not been clearly established yet. In this study, we investigated *IMPG2* expression and protein structure in the teleost zebrafish (*Danio rerio*), since it has a cone-dominant vision and thus, its retinal anatomy is quite similar to humans [8,29–31]. However, during evolution, the genome of teleost fishes underwent duplication. For this reason, many genes are found in two copies, named paralogues [32]. *IMPG2* is present as *impg2a* and *impg2b*. We obtained a phylogenetic tree of IMPG2 in different vertebrate species to investigate the extent of protein conservation during evolution. Moreover, since IMPG2 protein structure has largely been unstudied, we performed homology modelling of IMPG2 conserved domains both in human and in zebrafish. Finally, we analysed for the first time the expression of *impg2a* and *impg2b* in zebrafish, during early development and in the adult.

## Results

### IMPG2 sequence conservation analysis among vertebrates

Human IMPG2 is a 1241 residues protein with four topologically distinct regions: a signal peptide of 22 amino acids at the N-terminus, an extracellular topological domain (residues 23 to 1099), a helical transmembrane domain (residues 1100 to 1120) and a cytoplasmic topological domain (residues 1121 to 1241). It also contains two SEA domains and two EGF-like tandem repeats, together with 5 hyaluronan-binding motifs. The protein is also a target for post-translational modifications, such as glycosylation and phosphorylation, at different sites (UniProt database).

We first investigated IMPG2 conservation during evolution, by alignment of the protein sequence of each chosen species and subsequent generation of a phylogenetic tree. IMPG2 protein sequences of different species were retrieved from NCBI database, which indicated the presence of IMPG2 only in jawed vertebrates. We selected some of the most common species of different vertebrate groups to include in our analysis. We then used the Clustal Omega sequence alignment program to perform a multiple sequence alignment and generate a phylogenetic tree (Fig 1a), which reflects the distance in terms of sequence alignment between the different vertebrate species. The length of the branches is directly correlated with the difference between the sequences. For example, we observed that *Danio rerio* IMPG2a and IMPG2b and *Homo sapiens* IMPG2 protein sequences cluster separately. Such a sequence difference reflects the evolutionary distance between the two groups. Moreover, as reported in Section 1, the genome of teleost fishes underwent duplication [32], explaining the presence of two paralogues in *Danio rerio*, which cluster together in the phylogenetic tree (Fig 1a). Interestingly, the other teleost fishes included in our analysis (*Notobranchius furzeri* and *Oryzias latipes*) do not have paralogues. One explanation could be that the genomes of these two species underwent duplication, but the second copy of the gene lost its function during evolution and was no longer subjected to selective pressure. To understand in more detail the sequence conservation between the two zebrafish proteins and human IMPG2 we used UniProt database and we found some domains (SEA, EGF-like and transmembrane) that are conserved in all the three proteins (Fig 1b). By using the BLAST Alignment Tool, we demonstrated that both fish proteins share 65% identity with the region of the human protein where the conserved domains are located (residues 879-1238). These conserved domains were then deeper analysed by homology modelling, as described in the subsequent section.

**Fig 1:**
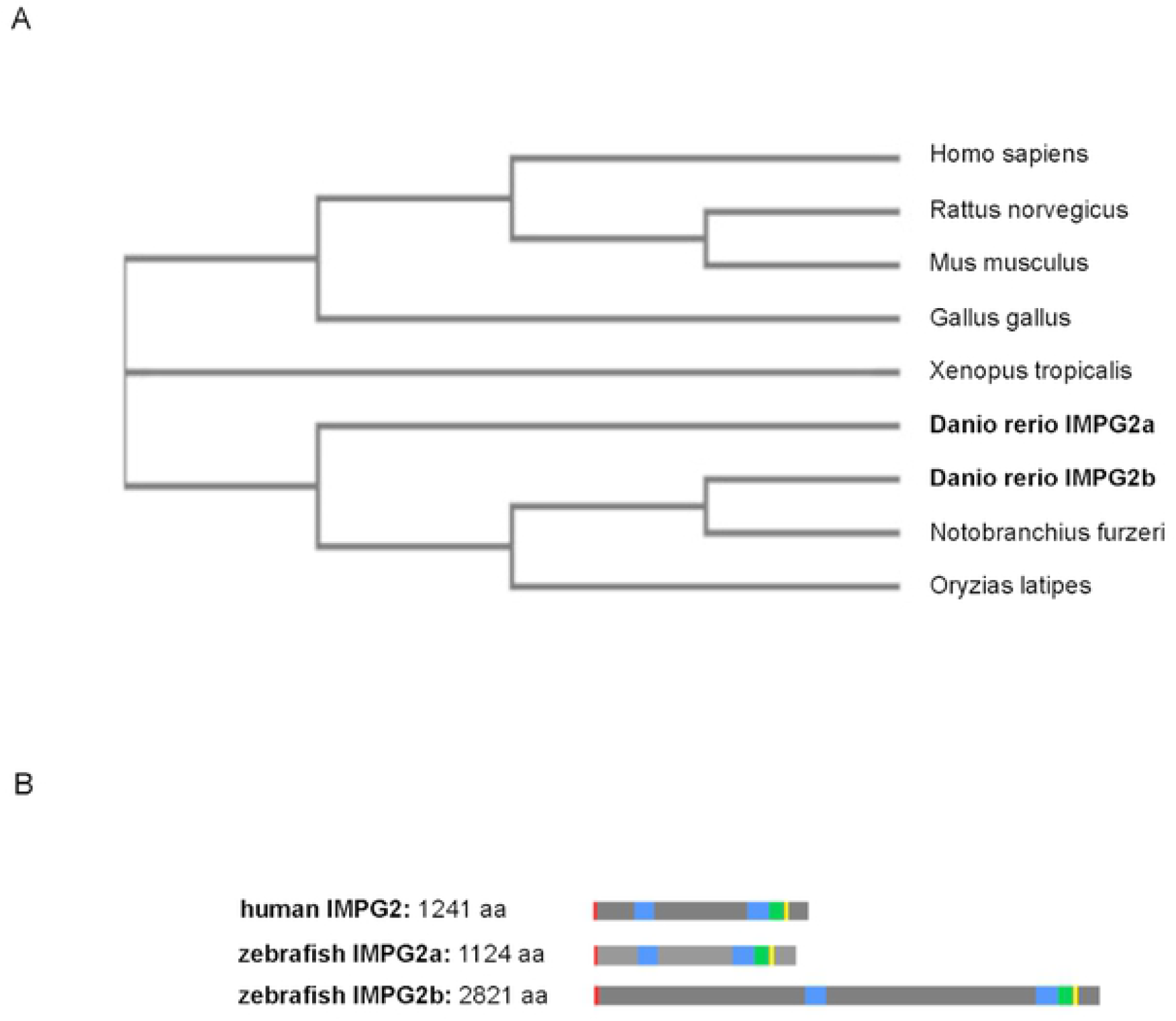
IMPG2 protein sequence conservation among vertebrates. (A) Protein sequence phylogenetic tree of IMPG2 among different vertebrate species obtained with the EMBL-EBI sequence analysis tool. Length of the branches reflects the distance between sequences. (B) Comparison of human IMPG2 protein and zebrafish IMPG2a and IMPG2b proteins. UniProt database was used to highlight the conserved domains in each of three proteins. In red, signal peptide; in light blue, SEA domain; in green, EGF-like domain; in yellow, transmembrane domain.

### Modelling of SEA and EGF-like domains in human IMPG2 and D.rerio IMPG2a and IMPG2b

Since little is known about the structure of the IMPG2 protein, we used iterative threading assembly refinement (iTasser) modelling to investigate the putative conformations of SEA and EGF-like conserved domains from their amino acid sequences [33]. The following sequences of the human protein were submitted to the iTasser webserver: 239-390, for SEA1, 896-1012 for SEA2 and 1012-1098 for the EGF-like tandem repeat. Domain identifications were obtained by checking Pfam [34] and Prosite [35] annotations. Then we used BLAST alignment to identify the sequences corresponding to the human domains in zebrafish IMPG2a and IMPG2b. The results are reported in Fig 2a. These sequences (also adding the non-overlapping residues at the terminals) were also submitted to the iTasser webserver for modelling. The best model proposed by iTasser for each domain was selected in terms of C-score. The predicted structures are represented in Fig 2b.

**Fig 2:**
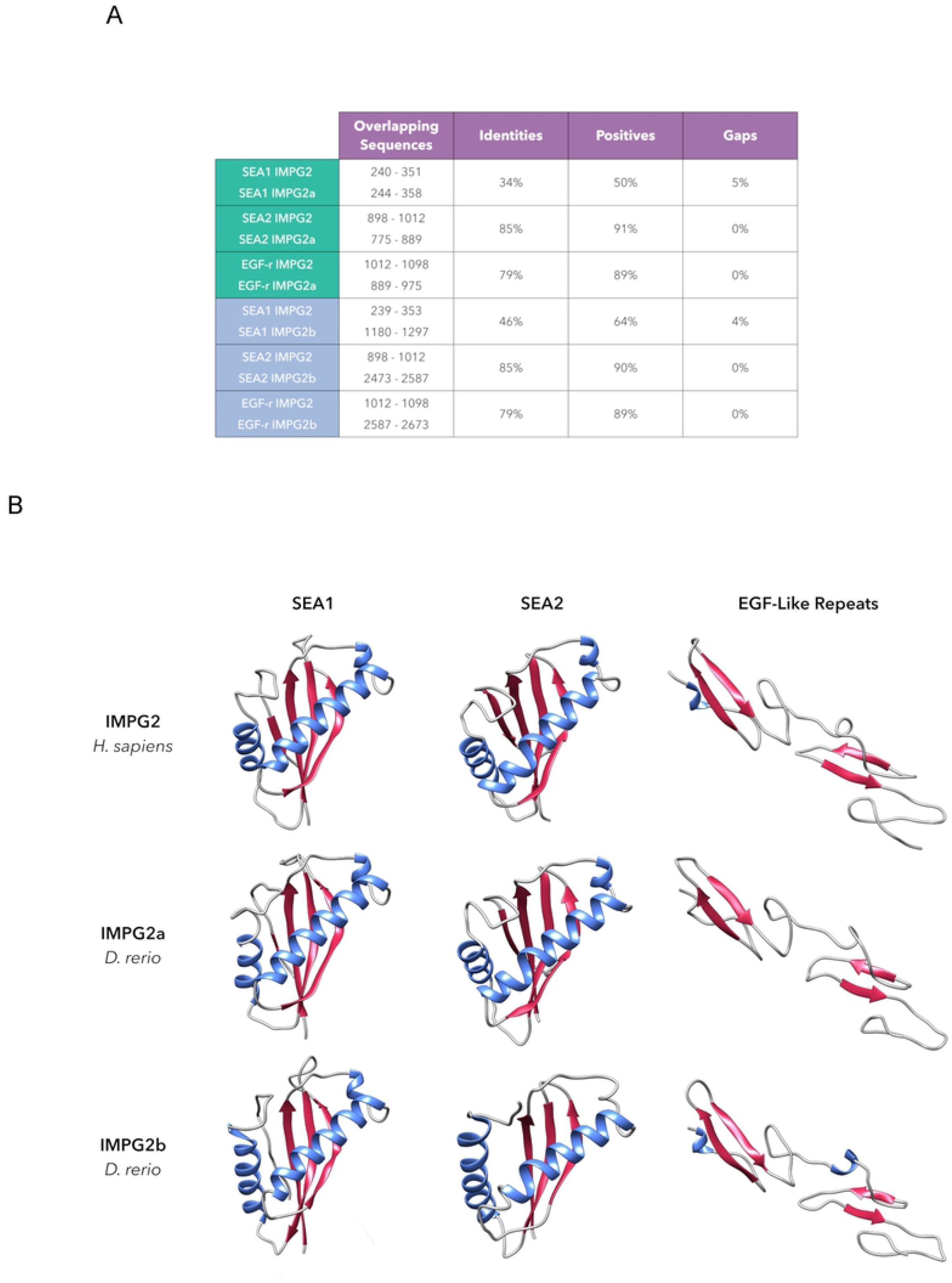
IMPG2, IMPG2a and IMPG2b conserved domains. (A) the figure shows SEA-1, SEA-2 and EGF-like domain sequences in zebrafish IMPG2a and IMPG2b with respect to the human protein. (B) Ribbon diagrams of structural models generated by iTasser of human IMPG2-SEA1, IMPG2-SEA2 and IMPG2-EGF-like-repeats, zebrafish IMPG2a-SEA1, IMPG2a-SEA2, and IMPG2a-EGF-like-repeats, zebrafish IMPG2b-SEA1, IMPG2b-SEA2 and IMPG2b-EGF-like-repeats. Secondary structures are depicted in different colours: blue, α-helices; red, β-strands; grey, coils.

### *impg2a* and *impg2b* mRNA expression

We investigated *impg2a* and *impg2b* mRNA expression in zebrafish during early embryonic development and in the adult fish. RT-qPCR experiments were performed on RNAs extracted from pools of whole embryos at different developmental stages and from pools of organs of adult fish. Results revealed very low expression of *impg2b* at 2.5 dpf and low *impg2a* mRNA levels at 3 dpf. However, *impg2b* and *impg2a* mRNAs start being significantly expressed at 3 dpf and 4 dpf, respectively (p<0.001, Tukey’s test following one-way ANOVA). In the analysis we compared the expression of *impg2a* and *impg2b* with that of *rhodopsin*, a strongly expressed photoreceptor-specific gene (Fig 3a). In the adult fish, RT-qPCR experiments showed that *impg2a* and *impg2b* are specifically expressed in the eye (Fig 3b). We next performed *in situ* hybridization (ISH) experiments on sections of embryos at 3, 5, and 7 dpf and adult fish, to investigate the localization of *impg2a* and *impg2b* mRNAs (Fig 3c). At 3 dpf the signals detected by ISH for both mRNAs are very low, becoming however detectable at 5 dpf with a specific expression in photoreceptor cell bodies. At 7 dpf the signal in the photoreceptor layer is stronger, and this high level of expression is maintained in the adult. Moreover, *impg2a* and *impg2b* expression seems to be found in both rods and cones, as already found in humans [36].

**Fig 3:**
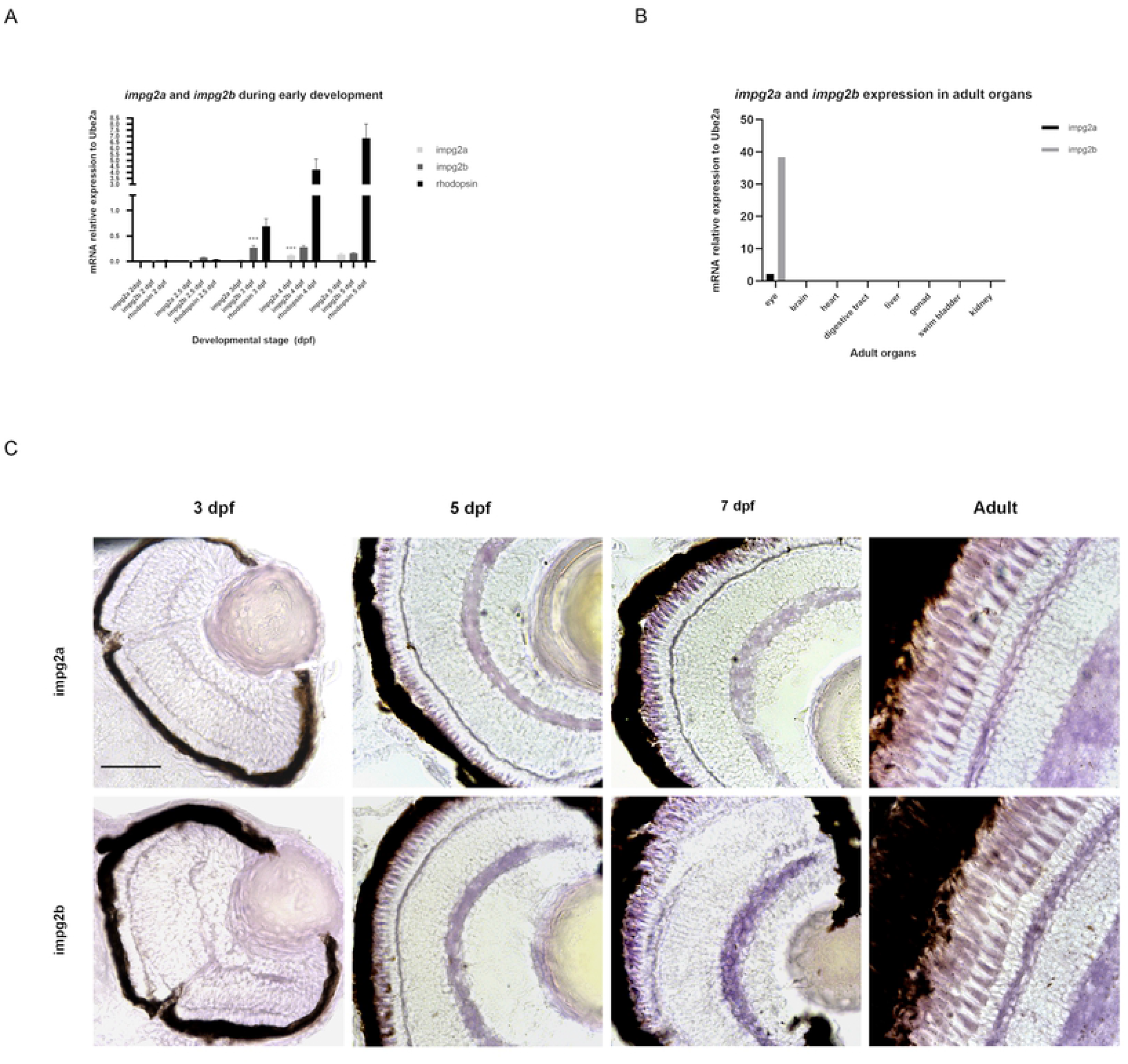
*impg2a* and *impg2b* expression during development and in the adult fish. (A) Quantitative RT-PCR analysis of *impg2a* and *impg2b* at different developmental stages. The expression of the two genes was compared to that of *rhodopsin; Ube2a* was used as control [37]. Data from three independent experiments revealed that *impg2a* starts to be significantly expressed at 4 dpf (p<0.001, Tukey’s test following one-way ANOVA, 2 dpf vs. 4 dpf, n=20 embryos per time point per experiment), whereas *impg2b* at 3 dpf (p<0.001, Tukey’s test following one-way ANOVA, 2 dpf vs. 3 dpf, n=20 embryos per time point per experiment). (B) mRNA expression levels of *impg2a* and *impg2b*, as obtained by RT-qPCR performed on pools of adult organs for each experiment. Data from three independent experiments showed specific expression of the two genes in the eye. (C) *In situ* hybridization experiments showing specific expression of both *impg2a* and *impg2b* mRNAs in the photoreceptor layer (black arrows) at different developmental stages and in the adult fish. Scale bars: 100 µm.

### IMPG2a and IMPG2b protein expression analysis during development and in the adult

To study protein expression during development and in the adult fish, we performed western blot experiments on pools of embryos at different developmental stages and pools of brains and eyes of adult fishes. We used a human IMPG2-specific antibody that recognizes both IMPG2a and IMPG2b proteins in zebrafish. The expression of the two proteins becomes detectable by western blot at 3 dpf (Fig 4a). Rhodopsin, a protein involved in the phototransduction cascade and expressed only in the outer segment of rod photoreceptor cells [38], also starts being expressed at 3 dpf. This suggests that IMPG2 expression accompanies photoreceptor maturation. Moreover, both proteins show a retina-specific expression, as observed for their mRNAs.

**Figure 4:**
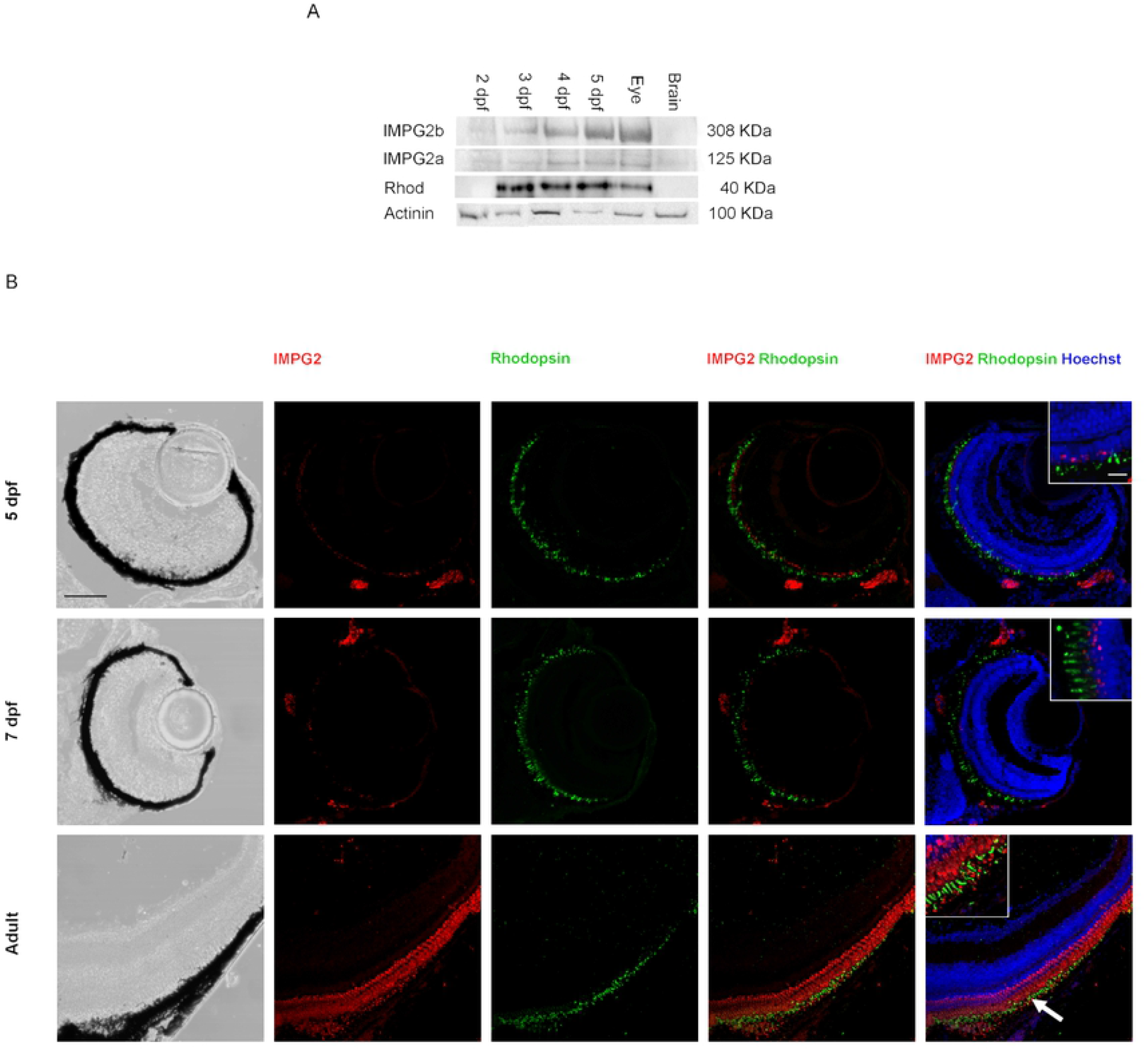
IMPG2a and IMPG2b localization during development and in the adult fish. (A) Representative western blot experiment showing expression of IMPG2a and IMPG2b starting from 3 dpf in zebrafish embryos and in the eye but not in the brain of adult zebrafish. Rhodopsin was used as a control. Actinin was used as housekeeping protein. (B) Immunohistochemistry for IMPG2 and Rhodopsin 1D4 performed on retina sections (14 µm) of zebrafish at 5 and 7 dpf and adult fish. IMPG2 signal starts to be detected at 5 dpf and becomes stronger in the adult retina, where it localizes in the photoreceptor cell body, in the photoreceptor outer segment (colocalizing with Rhodopsin, as indicated by the white arrow) and in the IPM. Brightfield images in the first column show the structure and the integrity of the retina and RPE. Scale bars: 50 µm. Inserts in merged images show higher magnification (scale bars: 100 µm).

To study the localization of IMPG2 in the zebrafish retina, we performed immunohistochemistry (IHC) experiments on sections of embryos at 5 dpf, 7 dpf and adult (Fig 4b). IMPG2 starts to be detected by IHC at 5 dpf and becomes stronger with development. It is particularly evident in the adult retina that IMPG2 localizes in the photoreceptor layer and in the interphotoreceptor matrix and it colocalizes with Rhodopsin, a rod oter segment (ROS) disk membrane specific marker [39], at the rod outer segments.

## Discussion

The extracellular matrix of the retina plays a key role in retinal function and disease [10; 40–42]. However, little is known about the function and the structure of many of its components, such as the proteoglycan IMPG2. Our study highlights the presence of SEA and EGF-like conserved domains in IMPG2 protein sequence of evolutionary distant vertebrate species such as H.sapiens and D.rerio. Interestingly, unlike other teleosts, D.rerio have two paralogues, IMPG2a and IMPG2b. This is not a peculiarity of IMPG2, indeed there are other genes characterized by having two paralogues in D.rerio but not in other teleost species, such as rs1a and rs1b. These two genes are orthologues of the human RS1 gene, whose mutation is associated X-linked retinoschisis (XLRS1 [MIM 312700]) in human [43]. Homology models of SEA and EGF-like conserved domains in human IMPG2 and zebrafish IMPG2a and IMPG2b show structure similarity of the domains in the two species. Importantly, we report for the first time the expression pattern of the two proteins in zebrafish, a valuable model organism for the study of human ophthalmological disorders. Unlike mouse models, they have cone-dominant vision like humans and the retina anatomy is similar to that found in humans [30,31]. Our experiments show expression of impg2a and impg2b mRNAs and proteins starting from 3 dpf. Moreover, we found both mRNAs and proteins to be specifically expressed in the photoreceptor layer and in the IPM. These data are consistent with the results of previous studies regarding localization of IMPG2 in rodents [1,27]. This work combines structural analysis of the conserved domains of human IMPG2 and zebrafish IMPG2a and IMPG2b, and expression analysis of impg2a and impg2b in zebrafish embryos and in the adult, providing novel insights into the biology of these disease-related genes.

## Materials and methods

### Animal care and maintenance

AB/TU wild-type zebrafish strain was used for all experimental procedures. Zebrafish were used under the approval of the OPBA of the University of Trento on Animal Welfare and Ministero della Salute (Project Number 151/2019-PR) and were raised following standard procedures [44].

### Phylogenetic tree

NCBI database was used to find orthologs to the human IMPG2 (*Homo sapiens* IMPG2, NP_057331.2; Norway ray IMPG2, XP_008766850.1; *Mus musculus* IMPG2, XP_017172459.1; *Gallus gallus* IMPG2, XP_015151604.1; *Xenopus tropicalis* IMPG2, XP_012813076.1; Danio rerio impg2a XP_017213311.1; *Danio rerio* impg2b XP_021329195.1; *Notobranchius furzeri* IMPG2, XP_015821571.1; *Oryzias latipes* IMPG2, XP_023806900.1). After choosing the vertebrate species to be included in the phylogenetic tree, IMPG2 protein sequences of these animals were obtained from both databases and a multiple protein sequence alignment was performed by using Clustal Omega sequence alignment program, provided by EMBL-EBI (https://www.ebi.ac.uk/Tools/msa/clustalo/). The same sequence analysis tool was used to generate a phylogenetic tree, based on protein sequence similarity.

### Modelling of SEA and EGF-like domains

Modelling of the SEA and EGF-Like domains was performed by using the iTasser webserver (https://zhanglab.ccmb.med.umich.edu/I-TASSER/). The submitted sequences for human IMPG2 were: 239-390 (SEA1), 896-1012 (SEA2) and 1012-1098 (EGF-like tandem repeat). The corresponding submitted sequences of IMPG2a were 239-371 (SEA1), 775-889 (SEA2) and 889-972 (EGF-like tandem repeat). The corresponding submitted sequences of IMPG2b were 1177-1333 (SEA1), 2473-2587 (SEA2) and 2587-2673 (EGF-like tandem repeat).

### RNA extraction and RT-qPCR

Total RNAs from pools of 15 embryos at different developmental stages and from pools of 3 adult eyes and 2 adult brains were extracted by Macherey Nagel NucleoSpin^®^ RNA. cDNA was synthesized by Super-Script^®^ VILO™ cDNA Synthesis Kit (Invitrogen). RT-qPCR was performed using KAPA SYBR^®^ FAST Master Mix (KAPA Biosystems) according to the manufacturer’s instructions. *Ube2a* was used as housekeeping gene and *rhodopsin* was used as reference gene, since it is highly expressed in the retina. Relative expression of each mRNA with respect to *Ube2a* mRNA was calculated as the average of three independent experiments. Expression analysis was performed using the CFX3Gene Manager (BioRad) software. Gene primers are listed in S1 Table.

## Protein extraction and western blot

Total proteins from pools of 15 embryos at different developmental stages and from pools of 3 adult eyes and 2 adult brains were extracted using RIPA buffer. 10 µg of total extract were resolved by SDS-PAGE, transferred to a nitrocellulose membrane and then incubated with the antibodies reported in S2 Table.

### Design of RNA probes

The genome browser Ensembl was used to find the exon sequences of the genes of interest. NCBI Primer-Blast (https://www.ncbi.nlm.nih.gov/tools/primer-blast/) was used to design the primers, following two criteria: ideal length between 20 and 24 bp and GC content between 42% and 52%. For the amplicon instead, a length between 400 and 1200 bp was chosen, with an ideal value of 600 bp. Finally, the T7 polymerase promoter sequence (GCGTAATACGACTCACTATAGGG) was added to the 5’ of the designed primers (S3 Table).

### Digoxigenin-labelled RNA probe synthesis

RNA extracted from a pool of 3 adult eyes with Macherey Nagel NucleoSpin^®^ RNA kit was reverse transcribed into cDNA, as described in section 4.4. The obtained cDNA was selectively amplified by using the primers in Table 2. PCR product was then purified using Wizard^®^ SV Gel and PCR Clean-Up System (Promega) and used for *in vitro* transcription with digoxigenin (DIG) labelled ribonucleotides and T7 polymerase. RNA samples were then treated with DNase I (Biolabs) to eliminate the cDNA templates and precipitated by adding salts (EDTA and 4M LiCl) and ethanol.

### ISH on retina sections

Embryos at different stages were fixed in 4% paraformaldehyde (PFA) at 4°C overnight, embedded in 30% sucrose at 4°C for 2-3 hours and included in OCT compound, with a head down position. Cryostat was used to obtain 14 µm retina sections. Briefly, sections were hybridized with 1 µg/ml probes overnight at 65 °C. The following day, saline sodium citrate (SSC) stringency washes and MABT (100mM maleic acid, 150mM NaCl, 0.1% Tween20) washes were performed. Sections were then incubated with blocking solution (1x MABT 1x, 2% Roche blocking reagent, 20% heat inactivated sheep serum) for 2 hours at RT and then with 1/2500 anti-DIG-AP antibody (Roche) in blocking solution overnight at 4 °C. The following day, after MABT washes, sections were coloured using NBT/BCIP (Roche).

### Immunohistochemistry

Embryos at different stages were fixed in 4% paraformaldehyde (PFA) at 4°C overnight, embedded in 30% sucrose at 4°C for 2-3 hours and included in OCT compound, with a head down position. Cryostat was used to obtain 14 µm retina sections. Immunohistochemistry was performed as follows: slides with sections were incubated in blocking solution (0.1% Triton X-100 and 0.5% BSA in 1× PBS) for 1 hour at room temperature (RT) and then incubated in diluted primary antibody in blocking solution (dilution specific for the primary antibody in use), at 4 °C overnight in humidified chamber. After 3 washes of 10 minutes in 1× PBS, 0.1% Triton X-100, slides were incubated in secondary antibody in blocking solution (1:1000), for 2 hours at RT in humidified chamber. Slides were then washed 3 times for 10 minutes 1× PBS, 0.1% Triton X-100 and then incubated with nuclear staining dye (1:10000; Hoechst 1) in 1× PBS for 10 minutes at RT. Following 3 washes of 10 minutes in 1× PBS, 0.1% Triton X-100, slides were mounted using Aqua-Poly/Mount coverslipping medium (Polysciences, Inc.).

Primary and secondary antibodies used for immunohistochemistry experiments are reported in S2 Table.

### Image acquisition

*In situ* hybridization images were taken on a Zeiss Axio Imager M2 up-right microscope using an EC Plan-Neofluar 40x/0.7 objective (Carl Zeiss Microscopy, LLC). Immunohistochemistry images were acquired using a Leica TCS SP8 confocal microscope equipped with an Andor iXon Ultra 888 monochromatic camera. The HC PL APO 40x/1.30 Oil CS2 (Leica Microsystems) objective was used for the acquisition. All figures were assembled in Fiji and Photoshop.

### Statistical analyses

All data are reported as mean ± SEM. Statistical analysis was performed using the GraphPad Software. Data groups from RT-qPCR experiments were compared by one-way ANOVA followed by Tukey’s test for multiple comparisons. Statistical significance level was set at p < 0.05. Values levels of statistical significance are described by asterisks (*p < 0.05; **p < 0.01; ***p < 0.001).

## Acknowledgments

We greatly thanks Ilaria Mazzeo (Model Organism Facility, Department of CIBIO, University of Trento) and Giorgina Scarduelli (Advanced Imaging Core Facility, Department of CIBIO, University of Trento) for their help in fish care and maintenance and image acquisition, respectively.

## Supporting Information

**S1 Table. Primers used for RT-qPCR experiments**.

**S2 Table. Antibodies used for Western blot (WB) and immunohistochemistry (IHC) experiments**.

**S3 Table. Primers used for Digoxigenin-labelled RNA probe synthesis**.

